# Combining multiple tools outperforms individual methods in gene set enrichment analyses

**DOI:** 10.1101/042580

**Authors:** Monther Alhamdoosh, Milica Ng, Nicholas J. Wilson, Julie M. Sheridan, Huy Huynh, Michael J. Wilson, Matthew E. Ritchie

**Affiliations:** CSL Limited, Bio21 Institute, 30 Flemington Road, Parkville, Victoria 3010, Australia.; ACRF Stem Cells and Cancer Division, The Walter and Eliza Hall Institute of Medical Research, 1G Royal Parade, Parkville, Victoria 3052, Australia.; Department of Medical Biology, The University of Melbourne, Parkville, Victoria 3010, Australia.; Molecular Medicine Division, The Walter and Eliza Hall Institute of Medical Research, 1G Royal Parade, Parkville, Victoria 3052, Australia.; School of Mathematics and Statistics, The University of Melbourne, Parkville, Victoria 3010, Australia.

## Abstract

Gene set enrichment (GSE) analysis allows researchers to efficiently extract biological insight from long lists of differentially expressed genes by interrogating them at a systems level. In recent years, there has been a proliferation of GSE analysis methods and hence it has become increasingly difficult for researchers to select an optimal GSE tool based on their particular data set. Moreover, the majority of GSE analysis methods do not allow researchers to simultaneously compare gene set level results between multiple experimental conditions.

**Results:** The ensemble of genes set enrichment analyses (EGSEA) is a method developed for RNA-sequencing data that combines results from twelve algorithms and calculates collective gene set scores to improve the biological relevance of the highest ranked gene sets. redEGSEA’s gene set database contains around 25,000 gene sets from sixteen collections. It has multiple visualization capabilities that allow researchers to view gene sets at various levels of granularity. EGSEA has been tested on simulated data and on a number of human and mouse data sets and, based on biologists' feedback, consistently outperforms the individual tools that have been combined. Our evaluation demonstrates the superiority of the ensemble approach for GSE analysis, and its utility to effectively and efficiently extrapolate biological functions and potential involvement in disease processes from lists of differentially regulated genes.

**Availability and Implementation:** EGSEA is available as an R package at http://www.bioconductor.org/packages/EGSEA/. The gene sets collections are available in the R package EGSEAdata from http://www.bioconductor.org/packages/EGSEA/.

## 1 Introduction

RNA-sequencing (RNA-seq) is a popular tool that enables researchers to profile the transcriptomes of samples of interest across multiple conditions in a high-throughput manner. The most common analysis applied to an RNA-seq dataset is to look for differentially expressed (DE) genes between experimental conditions. Gene set enrichment (GSE) often follows this basic analysis with the aim of increasing the interpretability of gene expression data by integrating *a priori* biological knowledge of the genes under study. This knowledge is usually presented in the form of groups of genes that are related to each other through biological functions and components, for example: genes active in the same cellular compartment, genes involved in the same signalling pathway or biological process, and so on. GSE methods calculate two statistics for a given dataset where pair-wise comparisons between two groups of samples, e.g. disease and control, are made: (i) a *gene-level* statistic calculated for each gene independently of other genes to identify DE genes in the dataset, and (ii) a *set-level* statistic derived for each gene set using the gene-level statistics (i) of its elements.

Statistical over-representation tests are the most commonly used methods for GSE analysis and are based on the top ranked DE genes obtained at a particular significance threshold. They suffer from a number of weaknesses, including the need to preselect the threshold and limited power on data sets with small numbers of DE genes. On the other hand, gene set tests, or so-called functional class scoring methods, do not assume a particular significance cut-off and also include the gene correlation in the calculation of the set-level statistics (Khatri et al., 2012). A third category of GSE methods incorporates the topology of the gene network in the significance statistics (Tarca et al., 2009). The definition of the null hypothesis in GSE analysis further categorizes these methods. *Competitive* tests assume the genes in a set do not have a stronger association with the experimental condition compared to randomly chosen genes outside the set. A second class of methods test a *self-contained* null hypothesis that assumes the genes in a set do not have any association with the condition while ignoring genes outside the set. Selfcontained methods tend to detect more gene sets when run on a large collection of gene signatures due to their efficiency in detecting subtle expression changes (Goeman and BÜhlmann, 2007).

In practice, GSE is applied on a large collection of gene sets and ranks them based on their relevance to the conditions under study. Various significance scores are used to assign gene set ranks. Most gene set tests are not robust to changes in sample size, gene set size, experimental design and fold-change biases (Tarca et al., 2013; Maciejewski, 2014). Given the diversity of approaches taken by different GSE analysis methods, reliance on any one method across different types of RNA-seq experiments, that may vary in scale (from large disease studies to small-scale experiments), complexity (simple two group comparisons through to more complex experimental designs) and noise level (patient samples versus more controlled samples obtained from model organisms), is bound to be sub-optimal. This issue has been widely discussed in the field of machine learning and several ensembling approaches have been proposed over the last three decades (Alhamdoosh and Wang, 2014). Ensemble methods have been shown to outperform individual methods in a number of studies, for example, PANDORA integrates multiple analysis algorithms to find a more accurate list of DE genes (Moulos and Hatzis, 2015).

To overcome this uncertainty problem in gene set ranking we propose a new GSE method, Ensemble of Gene Set Enrichment Analyses (EGSEA), which utilizes the gene set ranking of multiple prominent GSE methods to produce a new ranking that is more biologically meaningful than the results from individual methods. EGSEA is demonstrated to be useful in carrying out downstream analysis on RNA-seq data. It generates a dynamic web-based report that displays the enrichment analysis results of all selected algorithms along with several ensemble scores. The gene sets can be ranked based on any of the individual or ensemble scores. EGSEA also provides powerful capabilities to visualize results at different levels of granularity. Comparative analysis is also featured in EGSEA, allowing gene sets to be identified across multiple experimental conditions. Finally, although EGSEA has mainly been developed to analyze RNA-seq data generated from human and mouse samples, it can be easily extended to other organisms.

The remainder of this paper is organised as follows: first we provide a brief review of existing GSE methods. Next we describe the EGSEA approach and implementation details, the gene signature collections that have been compiled and the data sets that EGSEA is demonstrated on. Finally, results are presented and future directions for the project are laid out.

### 1.1 A review of current GSE methods

As EGSEA combines multiple gene set testing algorithms, we begin with an overview of current GSE methods. Some technical aspects of these methods are highlighted, with an emphasis on their similarities and differences.

Over-representation analysis (ora) methods perform Fisher’s hypergeometric test on each gene set to examine the significance of the overlap between alist of DE genes and the elements from a reference list of genes (Tavazoie et al., 1999). The set of DE genes is obtained by applying cut-off thresholds of gene-specific scores (e.g. false discovery rates (FDRs) and/or fold-changes). However, these gene-specific scores are not used in the calculation of the gene set scores which can lead to a number of limitations (Khatri et al., 2012) (e.g. strongly and weakly expressed genes are considered equally). On the other hand, enrichment score-based methods use gene fold-changes or other test statistics to order the list of DE genes. A random walk is then used to find the maximum deviation from a reference value (usually 0) and calculate enrichment scores, as in the GSEA algorithm (Mootha et al., 2003; Subramanian et al., 2005). Sample-based permutation is then applied to estimate the significance of the gene set scores. These methods assume that gene sets related to the experimental condition are dominant at the top or bottom of the gene list. Variants on this approach that use the absolute values of gene scores to rank genes before performing a random walk have also been suggested (e.g. ssgsea Barbie et al. (2009)).

Other approaches tend to summarize the gene statistics for each set using global statistics and then test for significance using a permutation test, e.g. safe (Barry et al., 2005), Category (Jiang and Gentleman, 2007), zscore (Lee et al., 2008), gage (Luo et al., 2009) and padog (Tarca et al., 2012). Although permuting phenotype labels maintains the relationship between genes, it requires a large sample size in each experimental condition to accurately estimate the statistical significance. Alternatively, gene permutation can be used to lessen the effect of sample size in spite of its gene independence assumption (Subramanian et al., 2005). The camera method estimates the inter-gene correlation for each gene set and adjusts the gene set statistic for this effect (Wu and Smyth, 2012). Rotation can also be used to carry out gene set testing on small data sets, as in the roast and fry methods (Wu et al., 2010). The roast algorithm allows for gene-wise correlation and can be applied in any experimental design. Since it utilizes a Monte Carlo simulation technique, it can be quite slow when run on a large collection of gene sets. Fry is a fast approximation that assumes equal gene-wise variances across samples, producing similar *p*-values to a roast analysis run with an infinite number of rotations.

Gene set statistics can be estimated in a variety of ways using simple statistics (e.g. the mean or sum of the statistics across all genes in a set) or more complicated approaches. Linear models are widely used for this purpose (Smyth, 2004), as in globaltest (Goeman et al., 2004), camera (Wu and Smyth, 2012), fry and roast (Wu et al., 2010), and allow multiple covariates to be included in the analysis. Several methods quantify gene set scores for each sample independently rather than for each experimental condition and then incorporate these scores into complex linear models to estimate the significance of a gene set in an experimental comparison. In other words, the gene expression data is transformed from the gene space into the gene set space. For example, the plage algorithm uses singular value decomposition (SVD) of the expression matrix for a set of genes to calculate pathway scores (Tomfohr et al., 2005). Similarly, gsva calculates a Kolmogorov-Smirnov-like rank statistic for every gene set in each sample and uses linear modelling to estimate the gene set significance for each experimental condition (Hänzelmann et al., 2013).

A relatively new trend that has emerged in GSE analysis incorporates the topology of the gene set (i.e. the interactions between gene products) into the gene set scoring functions and significance tests, e.g. SPIA (Tarca et al., 2009). It has recently been shown by Bayerlová et al. (2015) that such methods do not always outperform simple gene set testing methods. Namely, when a particular group of genes appears in many of the gene sets tested, they are unlikely to be influential in the gene set significance test. Tarca et al. (2012) showed that results from padog can be improved by emphasizing the genes that appear in a smaller number of gene sets in the gene set test. All GSE methods mentioned above perform *p*-value adjustments to account for multiple hypothesis testing.

## 2 Materials and Methods

### 2.1 Ensemble of gene set enrichment analyses

By extending the concept of ensemble modelling into GSE analysis, we propose a new method that combines multiple GSE analyses in order to generate a robust gene set ranking that offers an improvement over the ranking obtained by individual methods. EGSEA, an acronym for *Ensemble of Gene Set Enrichment Analyses*, utilizes the analysis results of twelve prominent GSE algorithms in the literature to calculate collective significance scores for each gene set. These methods include: ora (Tavazoie et al., 1999), globaltest (Goeman et al., 2004), plage (Tomfohr et al., 2005), safe (Barry et al., 2005), zs-core (Lee et al., 2008), gage (Luo et al., 2009), ssgsea (Barbie et al., 2009), roast, fry (Wu et al., 2010), padog (Tarca et al., 2012), camera (Wu and Smyth, 2012) and gsva (Hänzelmann et al., 2013). The ora, gage, camera and gsva methods test a competitive null hypothesis while the remaining seven methods test a self-contained hypothesis. Conveniently, the algorithm proposed here is not limited to these eleven GSE methods and new GSE tests can be easily integrated into the framework. Figure 1 illustrates the general framework of EGSEA that can be seen as an extension of a popular RNA-seq analysis pipeline.

**Figure 1:**
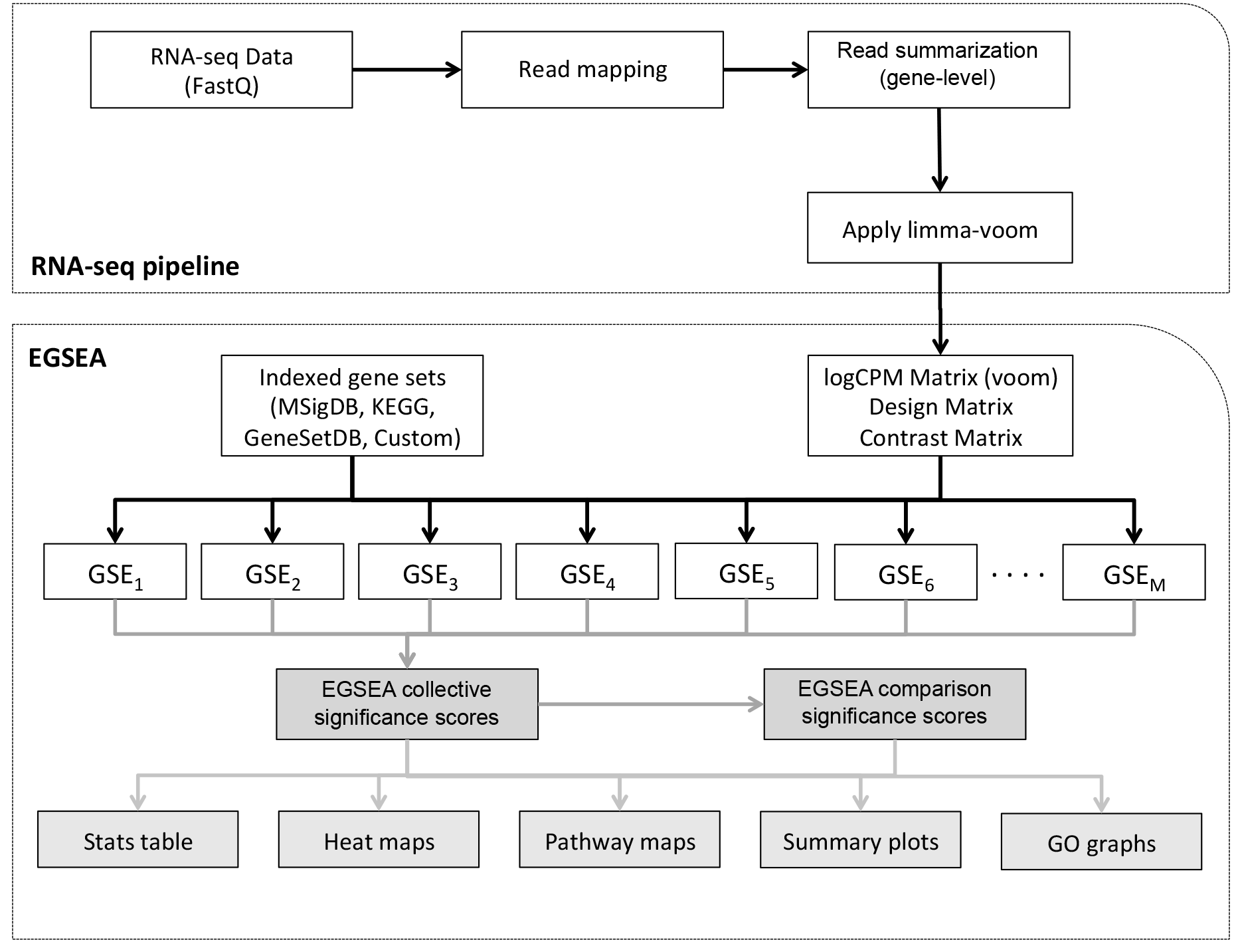
A schematic overview of the EGSEA pipeline for gene set enrichment analysis.

RNA-seq reads are first aligned to the reference genome and mapped reads are assigned to annotated genomic features to obtain a summarized count matrix. Most of the GSE methods were intrinsically designed to work with microarray expression values and not with RNA-seq counts, hence the limma-voom transformation is applied to the count matrix to generate an expression matrix (Law et al., 2014) applicable for use with these methods as has recently been shown (Rahmatallah et al., 2015). Since gene set tests are most commonly applied when two experimental conditions are compared, a design matrix and a contrast matrix are used to construct the experimental comparisons of interest. The target collection of gene sets is indexed so that the gene identifiers can be substituted with the indices of genes in the rows of the count matrix. The GSE analysis is then carried out by each of the selected methods independently and an FDR value is assigned to each gene set. Lastly, the ensemble functions are invoked to calculate collective significance scores for each gene set.

### 2.2 Problem formulation

Given an RNA-seq dataset *D* of samples from *N* experimental conditions, *K* annotated genes *g_k_(k = 1,…, K)*, *L* experimental comparisons of interest *C_l_(l = 1,…, L)*, a collection of gene sets Γ and *M* methods for gene set enrichment analysis, the objective of a GSE analysis is to find the most relevant gene sets in Γ which explain the biological processes and/or pathways that are perturbed in expression in individual comparisons and/or across multiple contrasts simultaneously. Numerous statistical gene set enrichment analysis methods have been proposed in the literature over the past decade. Each method has its own characteristics and assumptions on the analyzed dataset and gene sets tested. In principle, gene set tests calculate a statistic for each gene individually *f(g_k_)* and then integrate these significance scores in a framework to estimate a set significance score *h(γ_i_)*.

#### 2.2.1 Ensemble scoring functions

We propose seven statistics to combine the individual gene set statistics across multiple methods, and to rank and hence identify biologically relevant gene sets. Assume a collection of gene sets Γ, a given gene set γ_i_ ∈ Γ, and that the GSE analysis results of *M* methods on *γ_i_* for a specific comparison (represented by ranks 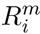 and statistical significance scores 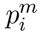, where *m* = 1,…, *M* and *i* = 1,…, |Γ|) are given. The ranks 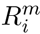 are calculated based on the order of *p*-values. When a tie occurs, the other test statistics of each individual method are used to break them. The EGSEA scores can then be devised, for each experimental comparison, as follows:

- The *p*-value score is the combination of *p*-values assigned to *γ_i_* and can be calculated in EGSEA using six different methods, which are described in Becker (1994) and Sutton et al. (2000), as follows:

1. Fisher’s method (FP) assumes that

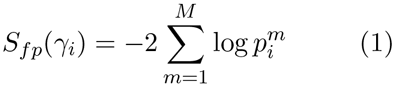
is a *Χ*^2^ distribution with 2*M* degrees of freedom (df).
2. The Logit method (LP) assumes that

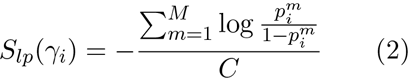

is a Student’s *t* distribution with *df* = 5*M*+4, where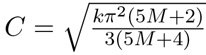.
3. The Summation of Z method (SZ) uses the weighted Z-test to calculate the combined *p*-value

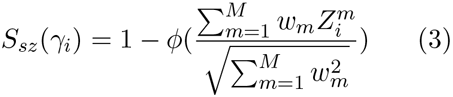

where 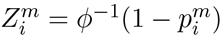, *w_m_* are weights, *ϕ* and *ϕ*^-1^ are the standard normal and its inverse. Equal weights are assigned for all base methods.
4. The Average method (MP) assumes that

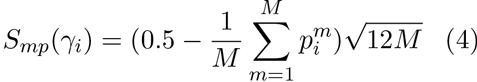

is a standard normal.
5. The Summation method (SP) sums the following series until the numerator becomes negative in order to estimate the combined *p*-value

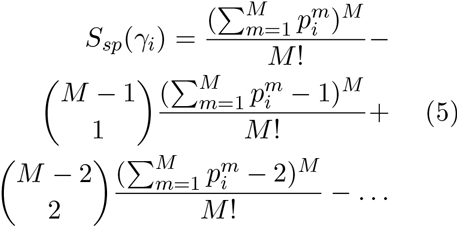
6. Wilkinson’s method (WP) calculates the probability of obtaining one or more significant *p*-values by chance in a group of *Mp*-values.

Note that the first three methods transform the *p*-values and then combine them. Finally, the Benjamini-Hochberg (BH) algorithm was applied to each *p*-value combining method (pCMs) to take into account the large number of tests being performed in parallel (Benjamini and Hochberg, 1995). It is worth noting that the p-value score assumes independence of the individual gene set tests, which is not a valid assumption here, hence they are not an accurate estimate of the ensemble gene set significance, but can still be useful for ranking results.

- The minimum *p*-value score is the smallest *p*- value calculated for *γ_i_*

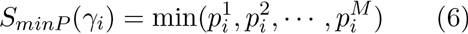

where 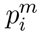 is the *p*-value calculated for the gene set *γ_i_* by the *m*-th GSE method
- The minimum rank score of *γ_i_* is the smallest rank assigned to *γ_i_*

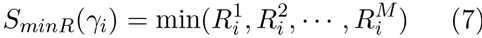

where 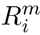 is the rank assigned by the *m*-th GSE method to the gene set *γ_i_*.
- The average ranking score is the mean rank across the *M* ranks

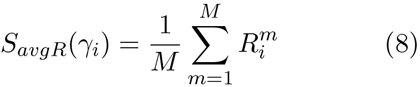
- The median ranking score is the median rank across the *M* ranks

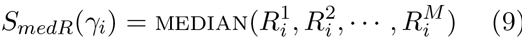

where MEDIAN is the classical median commonly used in statistics.
- The majority voting score is the most commonly assigned bin ranking

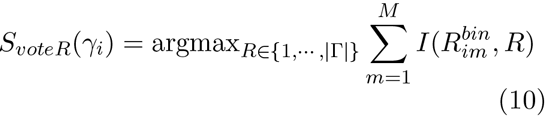

where 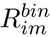 is the bin ranking of the gene set *γ_i_* that is assigned by the *m*-th method and is calculated using the following formula

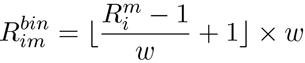

where *w* is the bin width. The bin ranking is used to obtain consensus ranking from multiple methods and thus a majority rank can be found.
- The significance score assigns high scores to the gene sets with strong fold-changes and high statistical significance

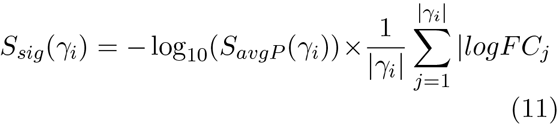
where *S_avgP_*(*γ_i_*) is the combined *p*-value and logFC_j_ is the log_2_ of the fold-change of the *j*-th gene in *γ_i_*. The significance score is scaled on the [0,100] range for each gene set collection.

#### 2.2.2 Comparative analysis

Unlike most GSE methods that calculate a gene set enrichment score for a given gene set under a single experimental contrast (e.g. disease vs. control), the comparative analysis proposed here allows researchers to estimate the significance of a gene set across multiple experimental contrasts. This analysis helps in the identification of biological processes that are perturbed by multiple experimental conditions simultaneously. For example, given three experimental conditions A, B and C, three pair-wise contrasts can be constructed (A versus B, A versus C and B versus C) and an EGSEA comparative analysis performed to find gene sets that are perturbed across two or three conditions simultaneously. Comparative significance scores are calculated for a gene set using Eqs. 1– 10 where the corresponding ensemble scores of individual pair-wise contrasts are substituted into these equations. In other words, the comparative ensemble scores for a given gene set *γ_i_* is calculated by replacing 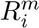 and 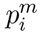 with the ensemble scores that are calculated for the *i*th experimental contrast.

An interesting application of the comparative analysis would be finding pathways or biological processes that are activated by a stimulation with a particular cytokine yet are completely inhibited when the cytokine’s receptor is blocked by an antagonist, revealing the functions uniquely associated with the signaling of that particular receptor as in the experiment below.

### 2.3 Gene set collections

The Molecular Signatures Database (MSigDB) (Subramanian et al., 2005) v5.0 was downloaded from http://www.broadinstitute.org/gsea/msigdb (05 July 2015, date last accessed) and the human gene sets were extracted for each collection (h, c1, c2, c3, c4, c5, c6, c7). Mouse orthologous gene sets of these MSigDB collections were adopted from http://bioinf.wehi.edu.au/software/MSigDB/index.html (Wu and Smyth, 2012). EGSEA uses Entrez Gene identifiers (Maglott et al., 2005) and alternate gene identifiers must be first converted into Entrez IDs. KEGG pathways (Kanehisa and Goto, 2000) for mouse and human were downloaded using the *gage* package. To extend the capabilities of EGSEA, a third database of gene sets was downloaded from the GeneSetDB (Araki et al., 2012) http://genesetdb.auckland.ac.nz/sourcedb.html project. In total, more than 25,000 gene sets have been collated and stored as R objects within the EGSEAdata package along with annotation information for each set (where available). Additional custom collections of gene sets can be easily added and tested using EGSEA. Supplementary Table 1 shows the number of gene sets in each collection and provides statistics on the set cardinalities and the overlap between gene sets. The Jaccard index is used to measure the similarity between two sets (Jaccard, 1912) and we calculate the third quar-tile and maximum overlap ratio between all possible pairs of gene sets in a collection. This analysis revealed that some collections contain many large gene sets. For example, the c2 collection from MSigDB contains 3,750 human gene sets with a median size of 37 and maximum size of 1,839. The Drug collection from GeneSetDB contains 7,032 human gene sets with a median size of 19. The overlap analysis shows that while some gene sets are very similar, the 3rd quartile of the Jaccard index is less than 2% for most of the collections.

### 2.4 Software implementation

EGSEA is implemented as an R package in the Bioconductor project (Gentleman et al., 2004) with parallel computation enabled using the *parallel* package. There are two levels of parallelism in EGSEA: (i) parallelism at the method-level and (ii) parallelism at the experimental contrast level. The results of an EGSEA analysis is stored in an object of S4 class named EGSEAResults. Several S4 generic methods were implemented to facilitate the integration of EGSEA in existing RNA-seq analysis pipelines as described in the software vignette (Alhamdoosh et al., 2016). A wrapper function was written for each individual GSE method to utilize existing R packages and create a universal interface for all methods. The ora method was implemented using the *phyper* function from the *stats* package, which estimates the hypergeometric distribution for a 2 × 2 contingency table. Statistical tests using *limma* were conducted in order to obtain the DE genes for ora. The implementation of roast, fry and camera was adopted from the *limma* package (Ritchie et al., 2015). Similarly, the GSE analysis methods of plage, zscore, gsva and ssgsea were available in the *gsva* package from Bioconductor. The gage, safe, globaltest and padog methods were implemented in the *gage, safe, global-test* and *padog* Bioconductor packages, respectively (Gentleman et al., 2004). EGSEA can be extended to include additional GSE methods through the implementation of new wrapper functions that the authors are happy to add on request. The *p*-value combining methods implementation was adapted from the *metap* package (Dewey, 2016).

Prior to running the EGSEA algorithm, an indexing mechanism is applied to the gene sets to transform gene identifiers into gene indexes that refer to the position of each gene in the count matrix. Finally, Jaccard coefficients were calculated for all possible pairs of gene sets using a parallel procedure with an exhaustive combinatorial calculation.

#### 2.4.1 Reporting capabilities of the software

Since the number of annotated gene set collections in public databases continuously increases and there is a growing trend towards generating dynamic analytical tools, our software tool was developed to enable users to interactively navigate through the analysis results by generating an HTML *EGSEA Report*. The report presents the results in different ways. For example, the *Stats table* displays the top *n* gene sets (where *n* is selected by the user) for each experimental comparison and includes all calculated statistics. Hyperlinks are enabled wherever possible, to access additional information on the gene sets such as annotation information. The gene expression fold-changes can be visualized using heat maps for individual gene sets or projected onto pathway maps where available (e.g. KEGG gene sets). The most significant Gene Ontology (GO) terms for each comparison can be viewed in a GO graph that shows their relationships. Similar reporting capabilities are also provided for the comparative analysis results of EGSEA.

Additionally, EGSEA creates summary plots for each gene set collection to visualize the overall statistical significance of gene sets. Two types of summary plots are generated: (i) a plot that emphasizes the gene regulation direction and the significance score (given in Eq. 11) of a gene set and (ii) a plot that emphasizes the set cardinality and its rank. EGSEA also generates a multidimensional scaling (MDS) plot that shows how various GSE methods rank a collection of gene sets. This plot gives insights into the similarity of different methods on a given dataset. Finally, the reporting capabilities of EGSEA can be used to extend any existing or newly developed GSE method by simply using only that method.

### 2.5 Simulated data

Simulated datasets were generated to evaluate the performance of EGSEA in various scenarios. First, a design matrix was defined for 5 case (Group 1) and 5 control (Group 0) samples, and a contrast matrix was created to compare Group 1 versus Group 0. In each simulation, expression matrices were generated with 15,000 genes of which 1,000 genes were selected to be DE and up-regulated and 1,000 genes were selected to be DE and down-regulated. The level of differential expression was defined in terms of log_2_ fold-changes so that the expression values of the DE genes were increased or decreased for the samples of Group 1 only by a particular amount.

To achieve this, log_2_ fold-changes were assumed to be normally distributed with mean 0 and gene-wise variances coming from a scaled-inverse chi squared distribution with 4 degrees of freedom. For the DE genes, the mean in Group 1 was systematically varied either up or down by a particular amount (between log_2_(1.3) and log_2_(2.3)) in order to simulate changes that ranged from subtle (30%) thorough to large (2.3 fold) differences. A prior standard deviation of 0.3 was used, i.e., the standard deviation of the gene-wise expression levels was drawn from 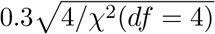. A total of 100 matrices were randomly generated for each simulation setting. A collection of 150 gene sets were generated such that 20 sets were composed of up-regulated genes only and a further 20 sets contained down-regulated genes only while the remaining sets were composed of non-DE genes. The gene sets were non-overlapping and the size was fixed to 50 genes for all sets.

### 2.6 Human IL-13 experiment

This experiment aims to identify the biological pathways and diseases associated with the cytokine Interleukin 13 (IL-13) using gene expression measured in peripheral blood mononuclear cells (PBMCs) obtained from 3 healthy donors. The expression profiles of *in vitro* IL-13 stimulation were generated using RNA-seq for 3 PBMC samples at 24 hours. The transcriptional profiles of PBMCs without IL-13 stimulation were also generated to be used as controls. Finally, an IL-13Rα1 antagonist (Redpath et al., 2013) was introduced into IL-13 stimulated PBMCs and the gene expression levels after 24h were profiled to examine the neutralization of IL-13 signaling by the antagonist. Single-end 100bp reads were obtained via RNA-seq from total RNA using a HiSeq 2000 Illumina sequencer. TopHat (Trapnell et al., 2009) was used to map the reads to the human reference genome (GRCh37.p10). HTSeq was then used to summarize reads into a gene-level count matrix (Anders et al., 2014). The TMM method (Robinson and Oshlack, 2010) from the *edgeR* package (Robinson et al., 2010) was used to normalize the RNA-seq counts. Data are available from the GEO database https://www.ncbi.nlm.nih.gov/geo/ as series GSE79027.

### 2.7 Mouse mammary cell experiment

Epithelial cells from the mammary glands of female virgin 8–10 week-old mice were sorted into three populations of basal, luminal progenitor (LP) and mature luminal (ML) cells as described in Sheridan et al. (2015). Three independent samples from each population were profiled via RNA-seq on total RNA using an Illumina HiSeq 2000 to generate 100bp single-end reads. The Subread aligner (Liao et al., 2013) was used to align these reads to the mouse reference genome (*mm10*) and mapped reads were summarized into gene-level counts using featureCounts (Liao et al., 2014) with default settings. The raw counts were normalized using the TMM method (Robinson and Oshlack, 2010). Data are available from the GEO database as series GSE63310. This dataset was first published in Sheridan et al. (2015), although no differential expression or GSE analysis was reported in this earlier study.

**Figure 2:**
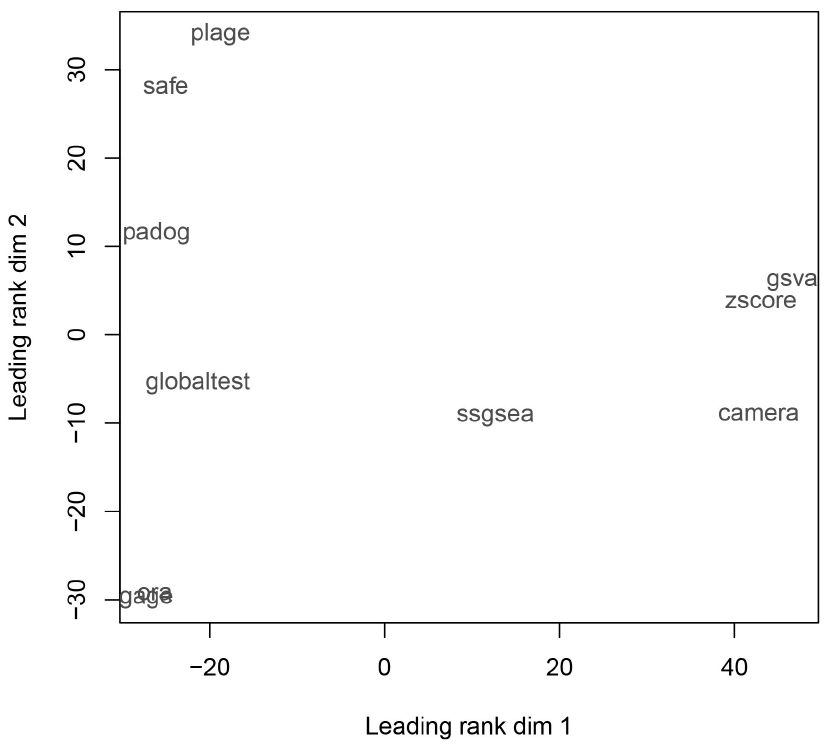
Multidimensional scaling plot based on the gene set rankings of the KEGG signalling and disease collections for ten GSE methods applied to the Human IL-13 vs. control dataset. Methods that perform similarly on this dataset cluster together.

## 3 Results and Discussion

The performance of the EGSEA method was evaluated using RNA-seq datasets that were either simulated or generated in the course of our research using either human or mouse samples (see Materials and Methods).

### 3.1 Performance on simulated data

To compare the performance of EGSEA and other methods in different settings, a cut-off threshold of 0.05 was used for the adjusted *p*-value in order to evaluate each algorithms’ retrieval power. Similarly, a cut-off threshold of 40 (top-ranked DE gene sets) was used to evaluate EGSEA’s vote, average and median ranking methods. The false discovery rate (FDR), true positive rate (Recall) and the F1-measure were calculated to measure the performance of EGSEA and the average over 100 simulated data sets of each configuration was reported along with the standard deviation. The F1-measure is the harmonic mean of recall and precision (1 - FDR). The performance indexes were calculated for each experiment using the six *p*-value combining methods (pCMs) and the EGSEA ranking scores. Eleven base methods, namely, camera, safe, gage, padog, plage, zscore, gsva, ssgsea, globaltest, ora and fry, were used in the following simulations unless otherwise stated.

First, the effect of the fold-change level on the performance of EGSEA was investigated. The level of differential expression simulated was varied between 1.3 and 2.3 fold and the performance indexes were calculated each time (Supplementary Table 3 and Table 1). As expected, the performance of EGSEA improves with increasing DE level, as most of the base methods tend to become more precise (Supplementary Table 4). At the lowest FC difference of 1.3, EGSEA gives an FDR as low as 1.61% and an F1-measure as high as 99.18% (both from Wilkinson’s method), and recall is 100% regardless of the pCM used (Table 1). EGSEA outperforms the majority of base methods at this FC level, with only two methods (safe and camera) performing slightly better in terms of their F1-measure (Supplementary Table 4). For simulated FC levels of 1.5 and 1.8, the true positive rate of 100% is maintained by EGSEA regardless of the pCM method while an FDR below 1% was obtained using the Average method (MP) and Summation methods (SP) at a FC of 1.8 (Table 1). For higher FC levels (≥ 1.5), while most of the individual GSE methods perform well, EGSEA is consistently amongst the top 4 methods (Supplementary Table 4). EGSEA generally controls the FDR for all of the pCMs except Fisher’s method which produces slightly more false positives (Table 1). The performance indexes from EGSEA’s ranking functions clearly show the advantage of using gene set rank rather than adjusted *p*-value when combining multiple GSE methods. The median rank is more robust than the vote and average ranks at low and high levels of simulated differential expression (Table 1). As the FC level increases, all EGSEA ranking scores achieve an F1-measure of 100%.

Second, the role of the number of base methods combined in EGSEA was investigated. Five experiments were designed for this purpose. The differential expression level was fixed at 1.3 in the five experiments and only the performance of EGSEA using Wilkinson’s method is shown here. The performance of EGSEA using the other pCMs is presented in Supplementary Table 2. The first experiment (E1) combined eleven methods: camera, safe, gage, padog, plage, zscore, gsva, ssgsea, globaltest, ora and fry, and aimed at highlighting the performance of EGSEA when all base methods are used. The second experiment (E2) excluded the ora method since it failed at retrieving any of the true positive gene sets. The third experiment (E3) excluded the worst performing methods (ora, gage and padog). The fourth experiment (E4) combined only the best five performing methods (safe, camera, fry, zscore, ssgsea). The fifth experiment (E5) included the best two methods of each style of test: the competitive methods camera and gsva and the self-contained methods zscore and fry. These simulation results clearly show that increasing the number of base methods benefits the ensemble performance even when a few weak methods are included in the ensemble (Table 1). Restricting EGSEA to the best performing methods, still gives FDR greater than those obtained from EGSEA based on all 11 methods (compare E1 with E4 and E5 in Table 2). Similarly, the performance of EGSEA drops only slightly when weak methods are removed (see E2 and E3 in Table 2). This observation is well addressed in the ensemble learning literature, where it has been shown that the performance of weak algorithms can be boosted dramatically by the majority (Freund, 1995).

### 3.2 Different methods produce different rankings

Our primary motivation was to improve the ranking of gene sets that are relevant to the experimental condition under study and thus improve the recall and precision of a GSE analysis. Various gene set tests assign different rankings to a collection of gene sets. To investigate this issue, the rankings assigned by ten GSE methods (camera, safe, gage, padog, plage, zscore, gsva, ssgsea, globaltest and ora) were obtained for the human IL-13 vs. control comparison. A multidimensional scaling (MDS) plot was generated using the ranks assigned by these ten methods to the 203 pathway maps in the KEGG signalling and disease collections. Fig. 2 clearly shows that some GSE methods perform more similarly on this particular collection and dataset than others. For example, camera, zscore and gsva seems to cluster together on the MDS plot. The Kendall rank correlation between zscore and gsva rankings was 0.62, between gage and ora was 0.56 and between camera and gsva was 0.49.

**Table 1:**
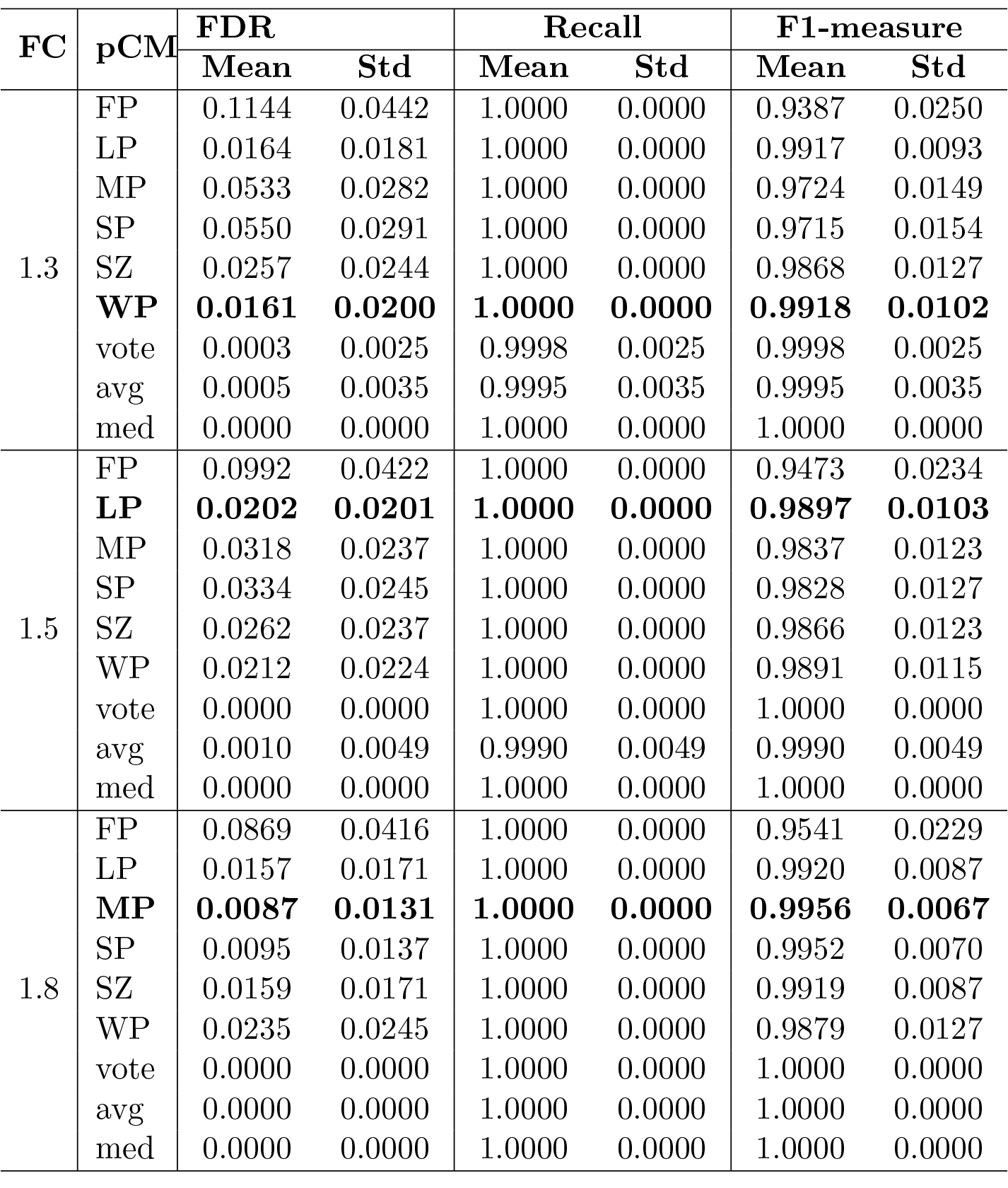
EGSEA’s performance at different levels of differential expression. FC is the differential expression level. FP, LP, MP, SP, SZ and WP stand for the Fisher, logitp, average, summation, summation of Z and Wilkinson p-value combining methods (pCMs), respectively. The best performing pCM is highlighted in bold for each FC configuration.

**Table 2:** EGSEA’s performance using a variable number of base methods with simulated FCs at the level of 1.3. Wilkinson’s method is used to combine p-values. The experiment E1 combines the eleven methods, E2 excludes ora, E3 excludes ora, gage and padog, E4 includes only camera, safe, zscore, ssgsea and fry, and E5 includes only camera, gsva, zscore and fry. The best performing configuration is highlighted in bold.

| ID | FDR |  | Recall |  | F1-measure |  |
| --- | --- | --- | --- | --- | --- | --- |
|  | Mean | Std | Mean | Std | Mean | Std |
| E1 | **0.0161** | **0.0200** | **1.0000** | **0.0000** | **0.9918** | **0.0102** |
| E2 | 0.0175 | 0.0210 | 1.0000 | 0.0000 | 0.9911 | 0.0108 |
| E3 | 0.0219 | 0.0231 | 1.0000 | 0.0000 | 0.9888 | 0.0119 |
| E4 | 0.0191 | 0.0221 | 1.0000 | 0.0000 | 0.9902 | 0.0114 |
| E5 | 0.0184 | 0.0212 | 1.0000 | 0.0000 | 0.9906 | 0.0109 |

Safe, padog and plage showed correlations with one and other of between 0.4 and 0.47 and globaltest and padog had a correlation of 0.42. Finally, the ranking produced by ssgsea was most dissimilar to the other methods, with correlations ranging between 0.12 and 0.32. Multidimensional scaling plots obtained using different gene set collections and data sets (Supplementary Figures S1-S9) were broadly similar, suggesting that the relationships between the different methods is consistent.

### 3.3 Performance on Human IL-13 experiment

Two experimental comparisons (IL-13 stimulated vs. control PBMCs, and IL-13R antagonist vs IL-13 stimulated PBMCs) were studied at the gene set level using EGSEA. Ten GSE methods, namely, camera, safe, gage, padog, plage, zscore, gsva, ssgsea, globaltest and ora, were used to calculate the collective EGSEA scores, and the *average rank* was used to identify significant gene sets. The vote rank was calculated using a bin width of 5. The analysis was conducted on the 203 signaling and disease KEGG pathways using a MacBook Pro machine that had a 2.8 GHz Intel Core i7 CPU and 16 GB of RAM. The execution time varied between 23.1 seconds (single thread) to 7.9 seconds (16 threads) when the HTML report generation was disabled. The execution time took 145.5 seconds when the report generation was enabled using 16 threads.

Table 3 shows the top ten pathways retrieved from the KEGG collections for these two experimental contrasts. Interestingly, the Asthma pathway was ranked as the first relevant pathway in the comparison between IL-13 stimulated PBMCs and control PBMCs. It has been shown that IL-13 is a key cytokine involved in the airway inflammation of patients with allergic asthma and IL-13 antagonists are successfully progressing through clinical development (Ingram and Kraft, 2012). It can be seen that the minimum ranking score assigned to Asthma by the ten GSE methods was nine and five methods assigned a rank higher than 13 to this pathway map. Supplementary Table 5 shows the ranks assigned by individual methods. The results also identified IL-13’s role in stimulating the intestinal immune network for IgA production (Cocks et al., 1993), which was retrieved as the second relevant pathway (Table 3). Although more than half of the testing GSE methods ranked this pathway higher than 25, EGSEA ranked it in the top 5 relevant pathways for IL-13 stimulated PBMCs. Similarly, Viral myocarditis disease appeared in the third position based on EGSEA ranking while most of the base GSE methods ranked it higher than 20. It has been found that IL-13 protects against myocarditis by modulating monocyte/macrophage populations (Cihakova et al., 2008). Moreover, the summary plot generated by EGSEA showed three pathways with very high significance scores (Fig. 3.A). They were the hematopoietic cell lineage signalling (hsa04640), the cytokine-cytokine receptor interaction (hsa04060) and the Staphylococcus aureus infection (hsa05150) pathways. The hsa04640 and hsa04060 were ranked 18*th* and 20*th* in the EGSEA results, respectively, while the hsa05150 pathway rank was higher than 20. The significance score *S_sig_* of these three pathways was greater than 80%. This means that these pathways are statistically significant and have a large number of DE genes for this contrast. It has been reported that the Staphylococcus aureus infection causes an increase in various cytokines including IL-13 (Wang et al., 2010).

**Table 3:** The top ten gene sets retrieved by EGSEA for the human PBMC data, based on the Average Rank scoring function. Two experimental contrasts were evaluated in this dataset. The gene set rank of the comparative analysis of these two contrasts is given in parentheses in the table below. The FDR is less than 0.05 for all sets.

| IL-13 Stimulated vs. Control |  |  |  |  | IL-13R Antagonist vs. IL-13 Stimulated |  |  |  |  |  |  |
| --- | --- | --- | --- | --- | --- | --- | --- | --- | --- | --- | --- |
| ID | Gene Set Name | Ranks |  |  |  | ID | Gene Set Name | Ranks |  |  |  |
|  |  | Vote | Avg. | Med. | Min. |  |  | Vote | Avg. | Med. | Min. |
| hsa05310 | Asthma (1) | 15 | 26.2 | 13 | 9 | hsa04621 | NOD-like recept. (6) | 20 | 29.3 | 25.5 | 9 |
| hsa04672 | Immune net. (8) | 20 | 31.6 | 26.5 | 12 | hsa04620 | Toll-like recept. (7) | 15 | 31 | 17 | 4 |
| hsa05416 | Viral myocarditis (2) | 20 | 32.4 | 20.5 | 7 | hsa05310 | Asthma (1) | 20 | 32.1 | 17.5 | 9 |
| hsa05146 | Amoebiasis (4) | 65 | 32.9 | 26.5 | 3 | hsa05414 | Dilated cardiomyo. (13) | 45 | 37.5 | 27.5 | 5 |
| hsa04961 | Calcium reabsorp. (11) | 10 | 36.5 | 25.5 | 4 | hsa05166 | HTLV-I infection (3) | 35 | 37.8 | 30.5 | 3 |
| hsa05134 | Legionellosis (5) | 10 | 40.7 | 35.5 | 8 | hsa05416 | Viral myocarditis (2) | 20 | 37.9 | 19 | 2 |
| hsa05166 | HTLV-I infection (3) | 15 | 41.9 | 32.5 | 9 | hsa05134 | Legionellosis (5) | 10 | 40.1 | 26 | 9 |
| hsa05020 | Prion diseases (>20) | 20 | 42.8 | 45.5 | 5 | hsa04623 | DNA-sensing (>20) | 55 | 40.6 | 40.5 | 8 |
| hsa05205 | Proteoglycans (17) | 50 | 46.2 | 44.5 | 15 | hsa05144 | Malaria (9) | 5 | 40.8 | 17.5 | 1 |
| hsa05145 | Toxoplasmosis (15) | 35 | 46.9 | 37.5 | 2 | hsa04064 | NF-kappa B sig. (>20) | 60 | 43.3 | 55.5 | 7 |

EGSEA analysis of the gene expression profiles comparing IL-13 stimulated PBMCs in the presence or absence of IL-13R antagonist retrieved Asthma as the third top pathway from the KEGG database (Table 3). Interestingly, only the CAMERA and ZSCORE methods ranked this pathway lower than 10 and its median rank across the ten methods was 17. This highlights the advantage of using an ensemble approach rather than relying on a single GSE method. The viral myocarditis pathway was ranked 6th for this contrast. The summary plot of IL-13R Antagonist vs IL-13 identified 4 gene sets: the cytokine-cytokine receptor interaction (hsa04060); Rheumatoid arthritis (hsa05323); Leishmaniasis (hsa05140) and; Staphylococcus aureus infection (hsa05150) with high significance score *S_sig_* (highlighted in blue in Fig. 3.A) that were not ranked in the top 10 gene sets (Table 3). Some of the base GSE methods assigned high rank to these pathways and therefore the *average rank* scores tend to be high. This shows the versatility of our proposed method, and also demonstrates how several ensemble scores can capture new knowledge about the investigated dataset. It is evident from the literature that IL-13 is increased in Rheumatoid arthritis serum (Tokayer et al., 2002) and plays a key role in the cutaneous Leishmaniasis (Hurdayal and Brombacher, 2014).

Finally, the EGSEA comparative analysis was performed on the two contrasts of this experiment, i.e., “IL-13 vs Control” and “IL-13R Antagonist vs IL-13”. This analysis retrieves gene sets that are perturbed in both contrasts and thus increases the power of gene

**Figure 3:**
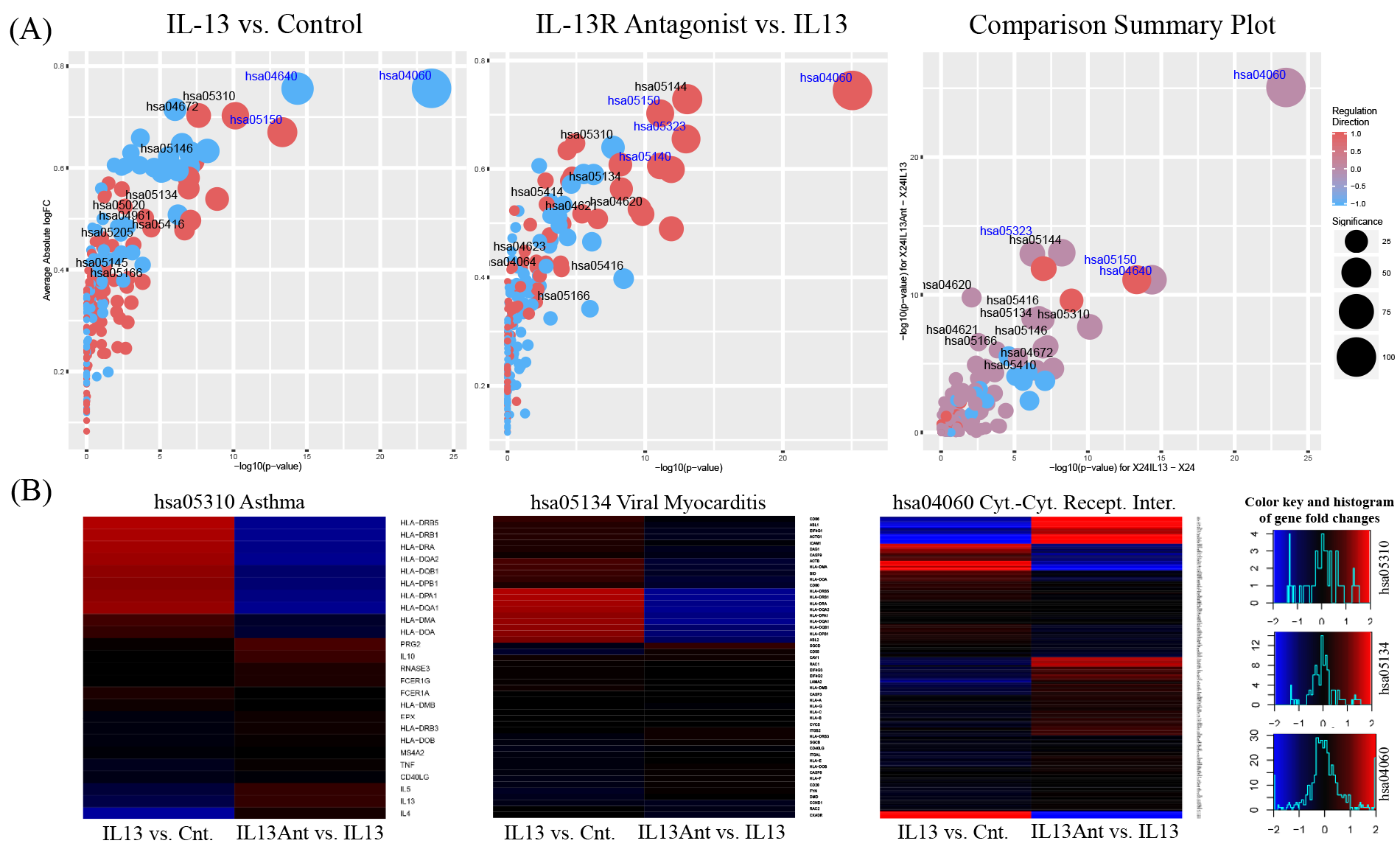
Visualization of the gene sets retrieved by EGSEA at diferent levels. (A) Summary plots of EGSEA on the human dataset. The IDs of the top ten pathways based on EGSEA average rank are highlighted in black font and the top five pathways based on EGSEA significance score whose average ranks are not in the top ten ranks are highlighted in blue font. The bubble size indicates the level of pathway significance. The red and blue colours indicate that the majority of gene set genes are up-or down-regulated, respectively. (B) Heat maps of the gene expression fold-changes in three selected gene sets.

set tests enabling an experiment-wide analysis. Here, the comparative analysis helped with investigating the neutralizing power of the IL-13R antagonist. Table 3 shows the rank of KEGG pathways (numbers in brackets) as assigned by the comparative analysis. The first two pathways discovered by this comparative analysis were Asthma and Viral myocarditis, respectively. Even though the cytokine-cytokine receptor interaction pathway did not appear in the top ten sets when IL-13 stimulated PBMCs were compared with control cells, it was assigned the twelfth rank in this analysis. The summary plot of the comparative analysis in Fig. 3.A shows KEGG gene sets coloured based on the average dominant regulation direction of genes and scaled based on the average significance score between the two contrasts. It is apparent that most of the pathways that were perturbed by IL-13 stimulation were inhibited by the IL-13R antagonist (coloured in purple). This gives an indication of the efficacy of the antagonist and highlights the utility of the comparative analysis. The Cytokine-cytokine receptor interaction (has04060), hematopoietic cell lineage signalling (hsa04640), Staphylococcus aureus infection (hsa05150) and Rheumatoid arthritis (hsa05323) pathways are all highly ranked (highlighted in blue). To further highlight the efficacy of the IL-13R Antagonist, Fig. 3.B displays heat maps of the fold-changes of gene expression in three exemplary pathways, namely, Asthma, Viral myocarditis and the cytokine-cytokine receptor interaction pathway. It can be clearly seen that the expression of individual genes is reversed in the different experimental conditions.

### 3.4 Performance on Mouse mammary cell experiment

Three experimental contrasts were studied from the mouse mammary cell experiment, i.e., basal versus luminal progenitor (LP) cells, basal versus mature luminal (ML) cells and ML versus LP. The *median rank* was used as a scoring function and a bin width of 5 was used for the vote ranking. Eight GSE methods were used as base methods for the EGSEA analysis: camera, safe, gage, padog, zscore, gsva, globaltest and ora. The analysis was conducted on the MSigDB c2 collection (of 4,722 gene sets) using the same machine that was mentioned earlier. The execution time varied between 182.1 seconds (single thread) to 72.9 seconds (16 threads) when the HTMLreport generation was disabled. The execution time took 147.5 seconds when the report generation was enabled using 16 threads.

In this experiment, the usefulness of the EGSEA comparative analysis was highlighted by analysing all three contrasts together. Table 4 shows the top ten gene sets retrieved from the c2 Curated Gene Set Collection of the MSigDB database. The LIM gene sets were generated previously by the same group on the same cell populations using microarrays (Lim et al., 2010) instead of RNA-seq. Five, out of six, of these earlier signatures, available in MSigDB were successfully retrieved by EGSEA using the RNA-seq data indicating that the current data is most similar to this earlier experiment, which is indeed the case. The average rank of the gene sets in Table 3 is relatively high which indicates that not all of the base GSE methods rank these signatures highly. We found that safe, padog and globabltest tend to assign very high ranks to the LIM gene sets, especially to the LIM Mammary Luminal Progenitor DN (M2576), which did not appear in this list of the top ten gene sets.

**Table 4:**
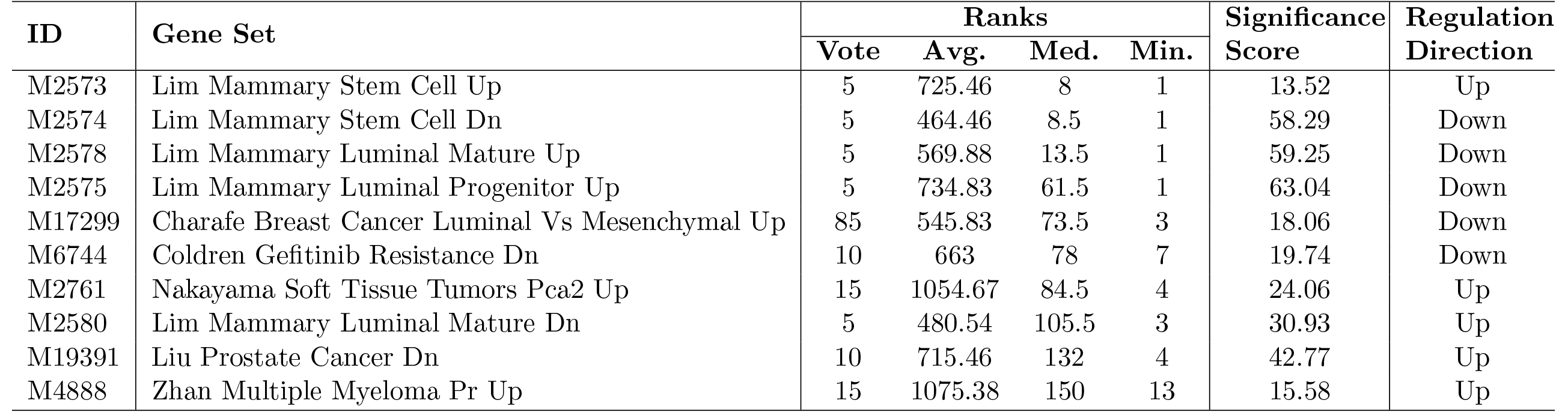
Comparative analysis results for three contrasts from the mouse mammary cell dataset.

## 4 Conclusion

Performing GSE analysis using a single method can be inefficient as determining which testing procedure is optimal for a given RNA-seq data set is a nontrivial task. Our results have shown that some methods may completely miss biologically meaningful associations in the data. To circumvent this problem, we developed a new approach, named EGSEA, that integrates multiple GSE tests into a single ensemble framework to improve the relevance of the biological processes identified for an experimental contrast. The analyses performed on RNA-seq datasets generated from human and mouse samples showed the advantage of our ensemble approach over using individual methods, with sensible results recovered in each example. EGSEA’s ability to perform a comparative analysis across multiple experimental contrasts simultaneously also helps overcome a limitation intrinsic to most GSE methods, which can only accommodate pair-wise comparisons one at a time.

EGSEA introduces an efficient solution to mine large databases of annotated gene sets. Our current implementation does not include topology-based GSE methods or support for microarray data, which we plan on adding in future releases of our software, along with interactive summary plots to enhance the user experience. Future research into the EGSEA approach will include an algorithm to select the appropriate number of methods to combine and the ability to assign variable weights to the different methods in a sensible way so that the results from less reliable GSE methods can be down-weighted in the analysis.

Since initiating this project, the Enrichment-Browser (EB) (Geistlinger et al., 2016) software, which takes a similar approach to EGSEA, has also been published. Compared to this approach, EGSEA combines twelve gene set testing methods and has been designed and tested specifically with RNA-seq data in mind, whereas EB combines four set-based methods and has been benchmarked primarily with microarray data. Our simulation results have shown that combining more methods is beneficial to the ensemble performance. Moreover, two of the four set-based methods (ora and safe) in EB fail when the expression signal is weak as shown in our simulations. An advantage of EB is that it includes four network-based methods, which as mentioned above we have yet to incorporate into EGSEA. Use of network-based methods is however limited to KEGG pathways at present and recent work by Bayerlová et al. (2015) has shown that network-based methods do not introduce a significant improvement on the retrieval performance relative to regular set-based methods that EGSEA currently focuses on. EGSEA also offers many more visualization options compared to EB. Finally, the various ensemble scores of EGSEA allow the ranking of gene sets in multiple ways to efficiently and effectively extract biological insights from large gene set collections.

## Acknowledgments

We thank Prof. Gordon Smyth, Dr Goknur Giner and Dr Aliaksei Holik from The Walter and Eliza Hall Institute of Medical Research (WEHI) for their critical feedback on this work, and Yifang Hu (WEHI) for generating R versions of many of the genes sets and Guido Pacini (WEHI) for providing simulation code.

